# Niche-FATE: niche-aware inference of cell-state transitions from spatiotemporal data

**DOI:** 10.64898/2026.11.24.768868

**Authors:** Junyao Jiang, Dong Won Kim

## Abstract

Cell-state transitions depend on intrinsic transcriptional programmes and local signals. Existing spatiotemporal mapping methods largely match cells by expression or position, not by received signals. Here we introduce Niche-FATE, an optimal transport framework integrating gene expression, spatial neighbourhoods and inferred ligand-receptor signalling to estimate transition probabilities and prioritise fate-associated signals. In simulations with changing tissue organisation, Niche-FATE improved fate inference and recovery of known fate-determining signals (mean AUPRC of 0.46 versus 0.29 for the best-performing alternative). In digit tip regeneration, Niche-FATE reduced inconsistent regenerative-to-scar transition assignments from 28% without niche information to 7%. Niche-FATE also improved developmental transition inference in mouse organogenesis. In two mouse models of Alzheimer’s disease, Niche-FATE linked predicted microglial transitions to local signalling, recovered amyloid plaque associations without plaque-location input and predicted transitions from disease-associated towards homeostatic-like states. Niche-FATE enables researchers to resolve divergent cellular outcomes and investigate their associated signalling environments.

## Introduction

Cell fate is shaped by both cell-intrinsic programmes and signals from the surrounding tissue. During development, cells with similar transcriptional states can adopt distinct fates in response to signals from neighbouring cells^1–4^. Similarly, during disease progression, local tissue environments can influence the emergence, maintenance and plasticity of cellular states^5–7^. Inferring these transitions from a cell’s transcriptional state alone may therefore overlook information contained within its local cellular and signalling environment^8^.

Spatial transcriptomics measures gene expression while preserving tissue organisation^9,10^. Spatial transcriptomic data sampled at multiple time points are increasingly available for development, regeneration and disease^11–13^. However, these measurements capture independent snapshots of tissue organisation rather than directly tracking individual cells over time. The computational challenge is therefore to infer cellular transitions between independently sampled time points.

Optimal transport-based methods, including WOT and TIGON, infer transitions between temporally sampled cell populations^14,15^. Spatial approaches, including spaTrack, moscot and STORIES, incorporate spatial organisation into the inference of cellular dynamics^16–20^. However, spatial geometry does not explicitly represent the signals individual cells receive. It may also become less informative when tissues undergo substantial reorganisation. Other approaches, including TopoVelo and CCCvelo, incorporate spatial dependencies or cell-cell communication into velocity-based inference^21,22^. Integrating the signals received by individual cells directly into temporal matching could help distinguish divergent cellular outcomes and identify candidate signals associated with these outcomes.

Here, we introduce Niche-FATE, a computational framework that integrates gene expression, spatial neighbourhoods and inferred received ligand-receptor signalling into unbalanced optimal transport. Niche-FATE estimates temporal cell-state transitions without requiring conserved tissue architecture and prioritises signals associated with alternative predicted outcomes. We benchmark Niche-FATE in simulations with known fate-determining signals and in mouse organogenesis. We then evaluate its ability to distinguish regenerative and scar-forming outcomes in digit tip regeneration and apply it to two mouse models of Alzheimer’s disease.

## Results

### Overview of Niche-FATE

Cells with similar transcriptional states can adopt distinct fates depending on their local cellular and signalling environments (Fig. 1a). Niche-FATE incorporates this principle into optimal transport-based inference of cell-state transitions. Rather than matching cells across time points based on their transcriptional states alone, Niche-FATE also considers their local cellular neighbourhoods and inferred received signals. Here, we refer to these two components collectively as the cellular niche.

**Fig. 1.**
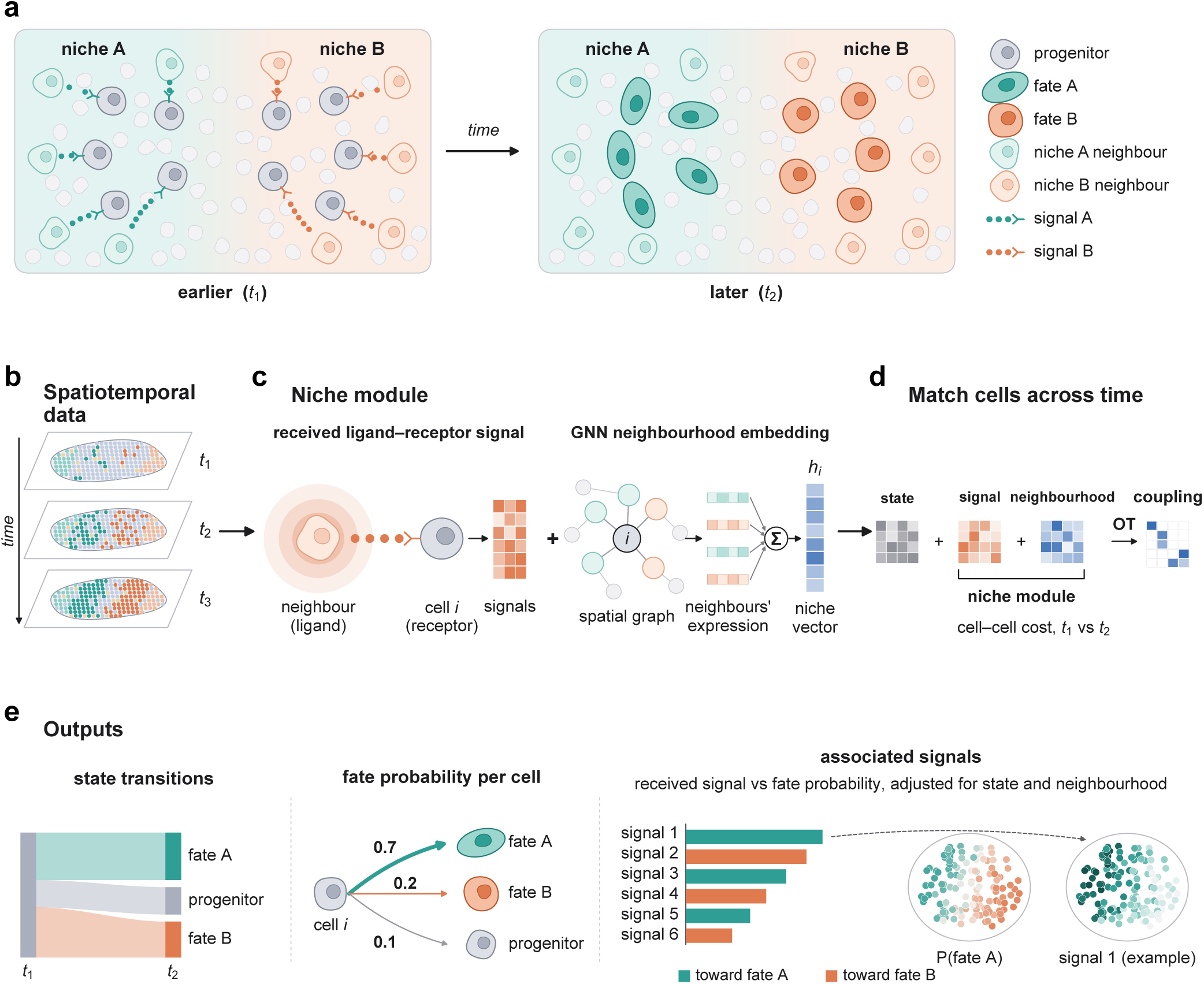
Overview of Niche-FATE. **a,** Progenitors with similar transcriptional states can adopt distinct fates in different cellular niches. **b,** Input spatial transcriptomic data sampled at multiple time points. **c,** Niche module comprising inferred received ligand-receptor signals and a GNN-based neighbourhood embedding. **d,** Cells are probabilistically coupled across consecutive time points using optimal transport based on transcriptional, signalling and neighbourhood dissimilarities. **e,** Outputs include cell-state transitions, cell-specific fate probabilities and signals associated with predicted cellular outcomes.

Niche-FATE takes spatial transcriptomic data sampled at two or more time points as input (Fig. 1b). For each cell, Niche-FATE characterises its niche using two complementary representations (Fig. 1c). The first is a received signalling profile, computed for each ligand-receptor pathway^23^ as the receptor expression of the focal cell multiplied by the distance-weighted ligand expression of its neighbours^24^. This provides an estimate of potential signalling based on ligand availability and receptor expression. The second is a neighbourhood embedding learned by a graph neural network, which captures the transcriptional states and spatial organisation of neighbouring cells. Cells across consecutive time points are then probabilistically coupled using unbalanced optimal transport, with a cost function combining transcriptional, signalling and neighbourhood dissimilarities (Fig. 1d). By incorporating niche information directly into the transport cost, Niche-FATE allows the local cellular environment to inform inferred transitions without requiring spatial alignment or conserved tissue architecture across time points. Under the benchmark conditions tested, Niche-FATE had the shortest runtime and lowest peak RAM usage among the compared methods (Extended Data Fig. 1).

Niche-FATE provides three outputs (Fig. 1e). State transitions describe the inferred relationships between cell states across time points. Fate probabilities quantify the fraction of each cell’s transported mass assigned to alternative target states and can be mapped onto the original tissue. Signal prioritisation identifies received signals associated with specific predicted outcomes. Niche-FATE computes partial correlations between received signals and fate probabilities while accounting for cellular state and neighbourhood information, with statistical significance assessed by spatial permutation (Methods). This analysis prioritises candidate niche signals associated with inferred transitions, providing testable hypotheses about how local signalling environments may influence cellular outcomes.

### Benchmarking fate inference and signal recovery in simulated tissues

To evaluate the performance of Niche-FATE, we generated simulated spatial time courses with known cell fates and fate-determining signals (Fig. 2a; Extended Data Fig. 2a; Methods)^25^. In each dataset, progenitors adopted one of two lineages according to their intrinsic transcriptional priming and received signals, and three of 20 ligand-receptor signals influenced lineage choice. We simulated two tissue configurations: shifting niches, in which spatial organisation changed between time points, and a fixed layout, in which tissue architecture was conserved. For each configuration, we simulated four signal strengths, ranging from no contribution to strong niche dependence. In the absence of niche effects, lineage choice was determined by intrinsic transcriptional priming and stochastic variation. When niche signals influenced lineage choice, progenitors with similar transcriptional states could adopt distinct fates depending on their local signalling environment (Fig. 2b).

**Fig. 2.**
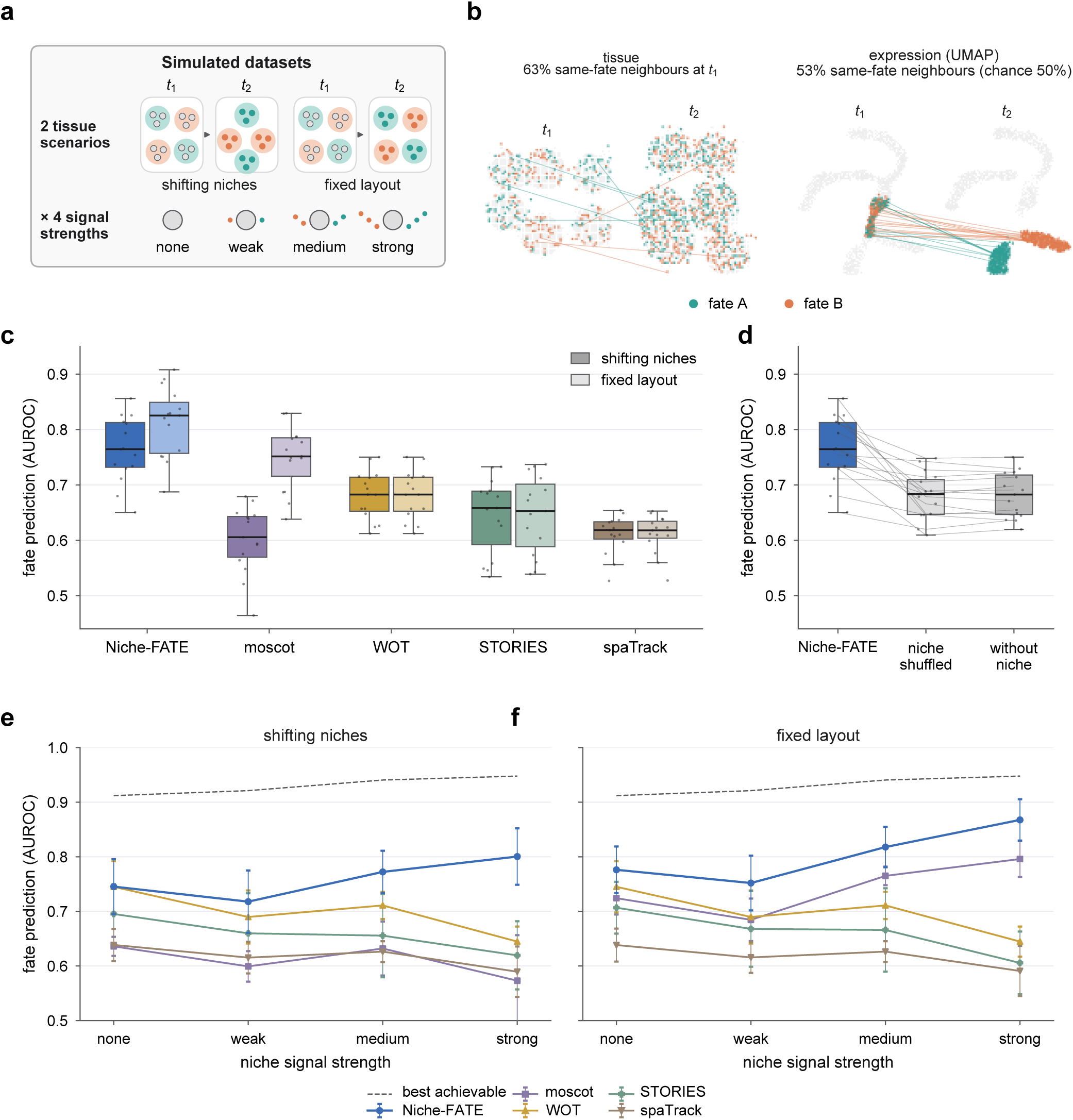
Niche-FATE improves fate inference in simulated tissues. **a,** Simulation design comprising two tissue configurations and four niche signal strengths. **b,** Representative simulated dataset showing progenitor fates in tissue and transcriptional space (UMAP). Percentages indicate the proportion of neighbouring progenitors sharing the same fate. **c,** Fate prediction performance, measured by the area under the receiver operating characteristic curve (AUROC), in tissues with shifting niches (dark) and a fixed layout (light). n = 15 simulated datasets per configuration. **d, A**blation analysis comparing Niche-FATE with spatially shuffled niche information and without the niche module in tissues with shifting niches. Lines connect results from the same simulated dataset. **e,f,** Fate prediction performance across four niche signal strengths in tissues with shifting niches **(e)** and a fixed layout **(f).** Data are presented as mean ± s.d. of five independent simulations. Dashed lines indicate the best achievable AUROC based on the simulated fate probabilities. In c,d, boxes indicate the median and interquartile range, and whiskers extend to 1.5 times the interquartile range.

We compared Niche-FATE with four existing methods: WOT, moscot, STORIES and spaTrack^14,16–18^. Across 15 datasets per tissue configuration, comprising 5 replicates at each of the 3 non-zero signal strengths, Niche-FATE achieved the highest mean AUROC (Fig. 2c). Niche-FATE achieved mean AUROCs of 0.76 in tissues with shifting niches and 0.81 in tissues with a fixed layout, compared with maxima of 0.68 and 0.75, respectively, among the other methods. Removing the niche module or constructing it from spatially shuffled cells reduced the mean AUROC from 0.76 to 0.68 in tissues with shifting niches (Fig. 2d), indicating that spatially organised niche information contributed to the improved inference.

We next examined how the contribution of niche signals to lineage choice affected inference accuracy (Fig. 2e,f). In the absence of niche effects, Niche-FATE achieved mean AUROCs of approximately 0.74 and 0.78 in the shifting and fixed configurations, respectively, reflecting the contribution of intrinsic transcriptional priming to lineage choice. As niche dependence increased from weak to strong, the AUROC of Niche-FATE increased from 0.72 to 0.80 in tissues with shifting niches and from 0.75 to 0.87 in tissues with a fixed layout. By contrast, the performance of expression-based WOT was lower under strong niche dependence. These results demonstrate that incorporating niche information improves fate inference when local signalling contributes to lineage choice, including when tissue organisation changes between time points.

We then examined whether incorporating niche information into fate inference also improved the recovery of fate-determining signals. For each method, we ranked the 20 ligand-receptor signals according to their associations with predicted fate probabilities after accounting for cellular transcriptional state (Methods). We evaluated the recovery of the 3 known fate-determining signals using the area under the precision-recall curve (AUPRC). In tissues with shifting niches, Niche-FATE achieved a mean AUPRC of 0.46, compared with a maximum of 0.29 among the other methods and 0.27 for random ranking (Extended Data Fig. 2b). In tissues with a fixed layout, Niche-FATE achieved a mean AUPRC of 0.52, compared with a maximum of 0.47 among the other methods evaluated (Extended Data Fig. 2b).

Incorporating niche information thus improved both fate inference and the recovery of known fate-determining signals in simulated tissues.

### Niche-FATE improves developmental transition inference and prioritises fate-associated niche signals

We next evaluated Niche-FATE using MOSTA, a Stereo-seq atlas of mouse organogenesis^11^. We analysed one sagittal section per developmental stage from E9.5 to E16.5, comprising approximately 500,000 spatial bins with annotated tissue identities (Fig. 3a). We inferred cell-state transitions between each pair of consecutive developmental stages. In the absence of lineage-tracing data, we evaluated the inferred transitions against previously curated developmental relationships between annotated cell types^17^. Transition accuracy was defined as the proportion of predicted transition probability assigned to developmentally consistent target populations, averaged across cell types.

**Fig. 3.**
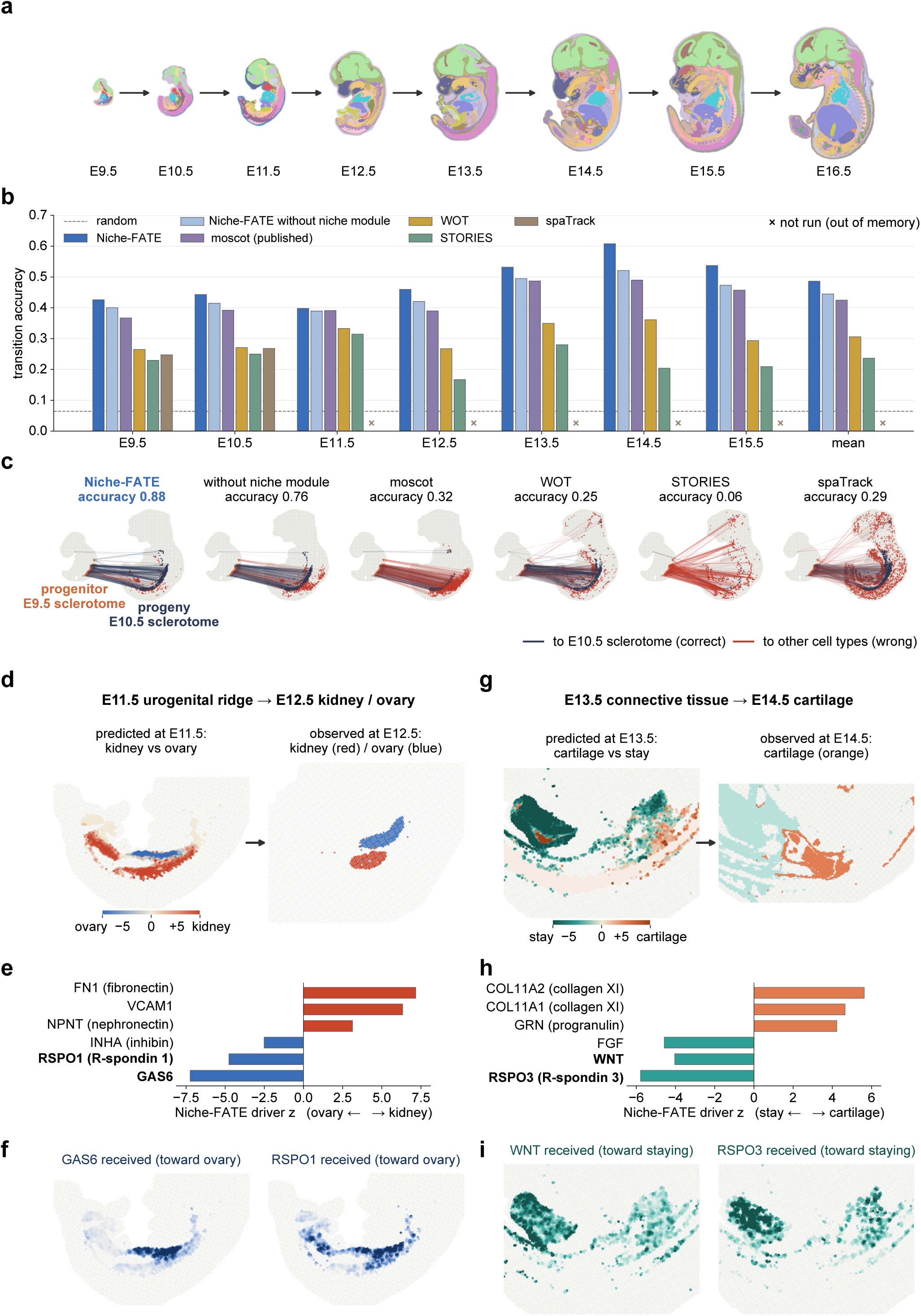
Niche-FATE improves transition inference and prioritises fate-associated signals during mouse organogenesis. **a,** Stereo-seq sections of mouse embryos from E9.5 to E16.5. **b,** Transition accuracy across consecutive developmental stages. Values for moscot were obtained from the original publication. Crosses indicate stage pairs for which SpaTrack could not be run owing to memory limitations. The dashed line indicates the random matching baseline. **c,** Inferred transitions from E9.5 sclerotome to E10.5 cell populations. Dark lines indicate transitions consistent with curated developmental relationships; red lines indicate inconsistent transitions. **d,** Predicted transitions from the E11.5 urogenital ridge towards kidney (red) or ovarian (blue) identity, alongside the observed tissue identities at E12.5. **e,** Niche signals associated with predicted kidney or ovarian outcomes. Positive and negative z-scores indicate associations with predicted kidney and ovarian outcomes, respectively. **f,** Spatial distributions of inferred GAS6 and RSPO1 reception. **g,** Predicted transitions from E13.5 connective tissue towards cartilage (orange) or retention of connective tissue identity (green), alongside observed cartilage at E14.5. **h,** Niche signals associated with predicted cartilage differentiation or retention of connective tissue identity. Positive and negative z-scores indicate associations with these respective outcomes. **i,** Spatial distributions of inferred WNT and RSPO3 reception.

Niche-FATE achieved the highest transition accuracy across all 7 developmental stage pairs, with a mean accuracy of 0.49, compared with 0.42 for moscot, 0.31 for WOT, 0.24 for STORIES and 0.06 for random matching (Fig. 3b). Removing the niche module reduced accuracy across all stage pairs, with the largest reduction observed between E14.5 and E15.5, from 0.61 to 0.52. Without niche information, 38% of pancreatic cells were mapped to the gut and 33% of spinal cord cells to the brain (Extended Data Fig. 3a,b). Incorporating the local tissue environment therefore improved the distinction between transcriptionally similar populations occupying different anatomical contexts. Similar improvements were observed at earlier developmental stages. For example, Niche-FATE assigned 88% of the transition probability from E9.5 sclerotome to E10.5 sclerotome, compared with 76% without the niche module, whereas moscot and STORIES predominantly mapped these cells to the adjacent spinal cord (Fig. 3c; Extended Data Fig. 3c).

Given the importance of intercellular signalling during organogenesis^26^, we next examined whether Niche-FATE could prioritise niche signals associated with distinct developmental outcomes. We first focused on the E11.5 urogenital ridge, which contains progenitors contributing to kidney and gonadal development^27^. Cells with higher predicted probabilities of transitioning towards kidney identity were predominantly located dorsally, adjacent to the sclerotome and dermomyotome, whereas those with higher predicted probabilities of transitioning towards ovarian identity occupied a more ventral domain facing the gut and liver (Fig. 3d; Extended Data Fig. 4a,b). This spatial distribution was consistent with the anatomy of the developing organs^27^. RSPO1 and GAS6 were associated with predicted ovarian outcomes (both P = 0.001; Fig. 3e, Supplementary Table 1), with higher inferred reception within the predicted ovarian domain (Fig. 3f). RSPO1 potentiates WNT signalling and is required for ovarian differentiation^28,29^. GAS6, the highest-ranked signal associated with predicted ovarian identity, is expressed in the embryonic gonad and supports primordial germ cell survival in culture^30^. Its association with predicted ovarian outcomes nominates GAS6 for further investigation in early gonadal development.

We next examined E13.5 connective tissue populations predicted to either acquire cartilage identity or retain connective tissue identity at E14.5. Cells with higher predicted probabilities of transitioning towards cartilage were predominantly located near existing cartilage primordia (Fig. 3g; Extended Data Fig. 4c,d), consistent with the recruitment of surrounding mesenchyme during cartilage development^31^. In contrast, reception of WNT and RSPO3 signals was associated with the predicted retention of connective tissue identity (both P = 0.001; Fig. 3h,i; Supplementary Table 1). Canonical WNT signalling can inhibit chondrogenic differentiation^32,33^. RSPO3, a WNT potentiator^34^ and the highest-ranked signal associated with connective tissue retention, emerges as a candidate regulator of non-chondrogenic mesenchymal identity.

### Niche-FATE distinguishes regenerative and scarring outcomes and prioritises regeneration-associated signals

We next evaluated Niche-FATE in a system with experimentally defined regenerative and scarring outcomes. Mice regenerate amputated digit tips following distal amputation that preserves the nail, whereas more proximal amputation that removes the nail results in fibrotic scarring (Fig. 4a)^35^. We analysed a Xenium spatial transcriptomic dataset comprising approximately 510,000 cells from regenerative digits at 5, 7, 10 and 14 days post amputation (dpa) and non-regenerative digits at 5, 7 and 14 dpa (Fig. 4b; Extended Data Figs 5 and 6a)^36^.

**Fig. 4.**
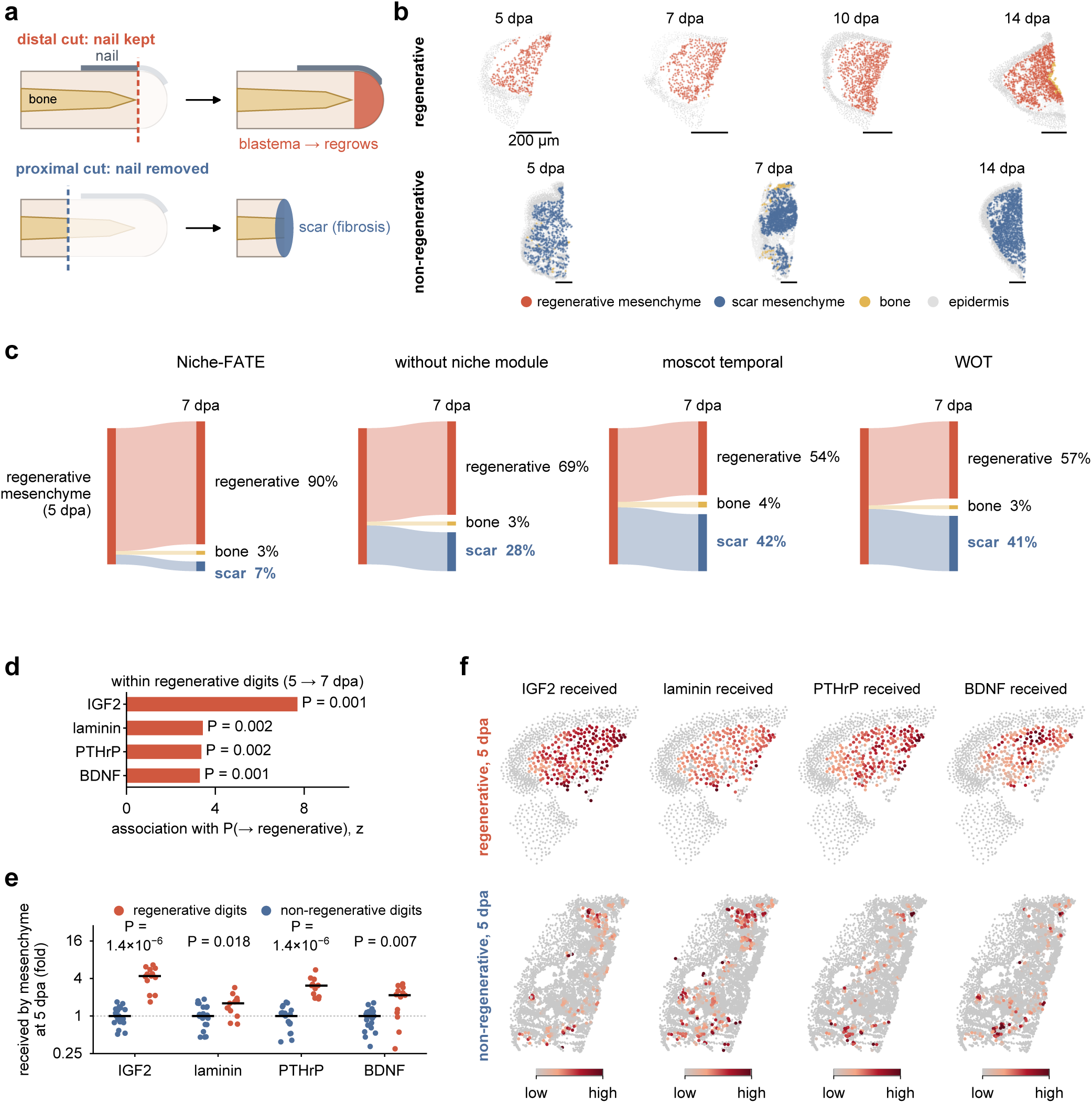
Niche-FATE distinguishes regenerative and scarring outcomes and prioritises regeneration-associated signals. **a,** Schematic of mouse digit tip amputation. Distal amputation preserves the nail organ and results in regeneration, whereas proximal amputation removes the nail organ and results in fibrotic scarring. **b,** Representative spatial transcriptomic sections of regenerative and non-regenerative digits at the indicated time points. dpa, days post-amputation. Scale bars, 200 µm. **c,** Inferred transitions from regenerative mesenchyme at 5 dpa to regenerative mesenchyme, bone or scar mesenchyme at 7 dpa using Niche-FATE, Niche-FATE without the niche module, moscot and WOT. **d,** Niche signals associated with the predicted retention of regenerative mesenchymal identity within regenerative digits, assessed using a spatial permutation test. **e,** Comparison of inferred signal reception by mesenchymal cells in regenerative and non-regenerative digits at 5 dpa. Each point represents one digit. Horizontal lines indicate medians. P values were calculated using a two-sided Mann-Whitney U test. **f,** Spatial distributions of the inferred signals shown in e in representative regenerative and non-regenerative digits at 5 dpa.

To assess whether niche information distinguishes these alternative outcomes, we pooled regenerative and non-regenerative digits and inferred cell-state transitions between 5 and 7 dpa. Niche-FATE assigned 90% of the transition probability from regenerative mesenchyme at 5 dpa to regenerative mesenchyme at 7 dpa and only 7% to scar mesenchyme (Fig. 4c). Removing the niche module increased the proportion assigned to scar mesenchyme to 28%, whereas moscot and WOT assigned 42% and 41%, respectively. Thus, incorporating niche information reduced transitions inconsistent with the known regenerative outcome, improving the distinction between mesenchymal populations associated with regeneration and scarring.

We next examined whether Niche-FATE could prioritise signals associated with the maintenance of regenerative mesenchyme. To distinguish these associations from general differences between regenerative and non-regenerative conditions, we restricted the signal prioritisation analysis to mesenchymal cells within regenerative digits. Cells with higher predicted probabilities of retaining regenerative mesenchymal identity showed higher inferred reception of IGF2 (P = 0.001), laminin (P = 0.002), PTHrP (P = 0.002) and BDNF (P = 0.001; Fig. 4d; Extended Data Fig. 6b; Supplementary Table 2). IGF signalling and laminins have established roles in tissue regeneration^37–41^, whereas PTHrP and BDNF are implicated in bone repair and skeletal muscle regeneration, respectively^42,43^.

We then asked whether the signals prioritised within regenerative digits differed between the experimentally defined regenerative and non-regenerative conditions. At 5 dpa, inferred reception of all 4 signals was higher in regenerative mesenchyme than in mesenchyme from non-regenerative digits (IGF2, P = 1.4 x 10^−6^; laminin, P = 0.018; PTHrP, P = 1.4 x 10^−6^; BDNF, P = 0.007; Fig. 4e,f). These findings nominate signals associated with both predicted mesenchymal fate and tissue regenerative capacity for further investigation in mammalian digit regeneration.

### Niche-FATE links local signalling environments to predicted microglial state transitions in Alzheimer’s disease

Finally, we applied Niche-FATE to neurodegeneration. In Alzheimer’s disease (AD), microglia near amyloid plaques transition from a homeostatic state towards a disease-associated microglial (DAM) state, in part through TREM2 signalling^44,45^. We analysed a public Xenium dataset of TgCRND8 mice, an amyloid model, at 2.5, 5.7 and 17.9 months, together with wild-type controls^46^. Amyloid plaques were imaged in the 17.9-month TgCRND8 section (Fig. 5a), and microglial states were annotated based on established marker genes (Extended Data Fig. 7a,b).

**Fig. 5.**
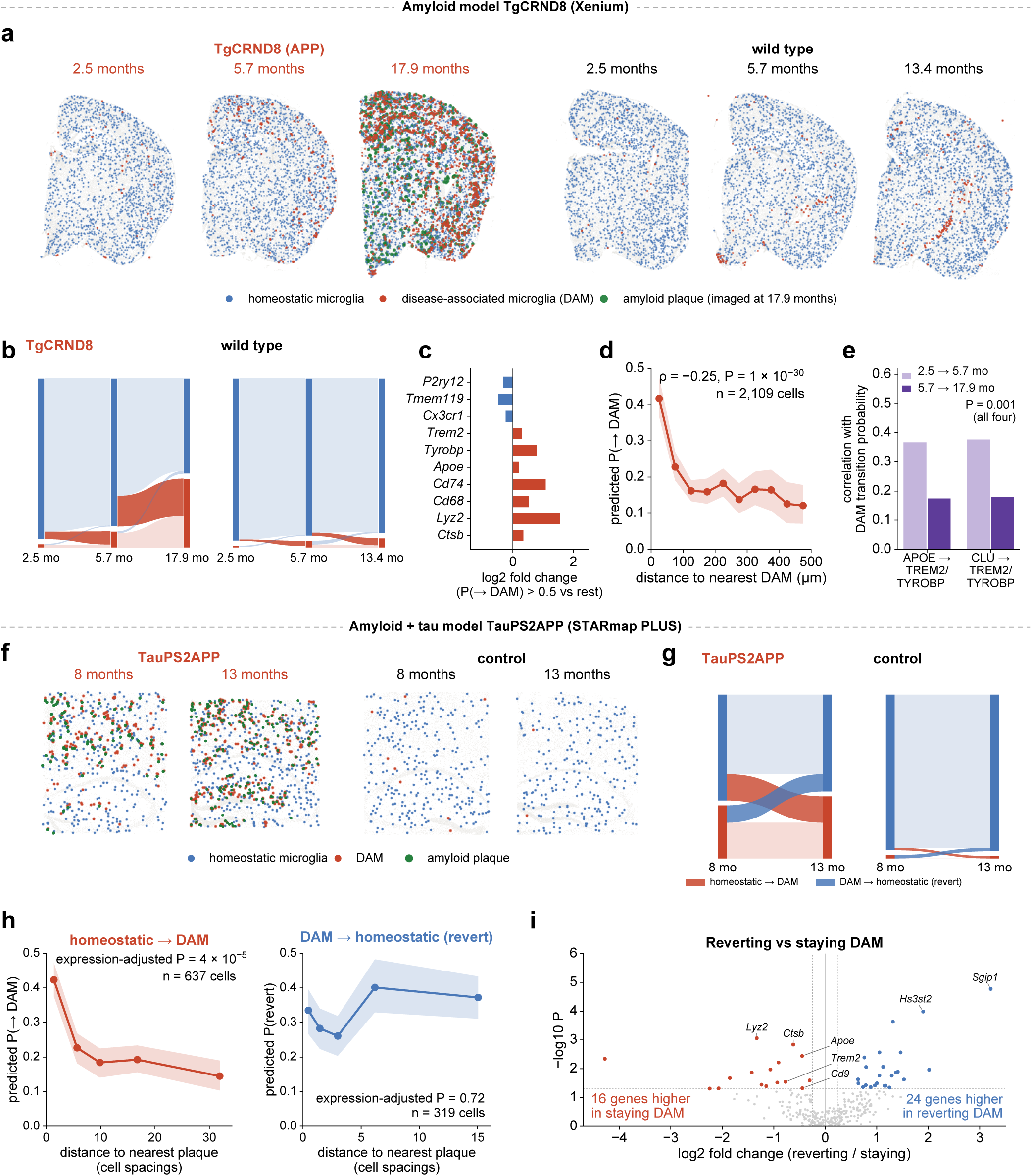
Niche-FATE predicts microglial state transitions and identifies fate-associated niche signals in models of Alzheimer’s disease. **a,** Spatial distributions of homeostatic and disease-associated microglia (DAM) in TgCRND8 and wild-type mouse brain sections profiled using Xenium. Amyloid plaques were imaged in the 17.9-month TgCRND8 section. b, Inferred transitions between homeostatic microglia and DAM in TgCRND8 and wild-type mice. Blue and red indicate homeostatic microglia and DAM, respectively. c, Log_2_ fold changes in marker gene expression between homeostatic microglia with a predicted probability of acquiring DAM identity greater than 0.5 and the remaining homeostatic microglia. d, Association between predicted DAM acquisition probability and distance to the nearest DAM (Spearman’s rank correlation; n = 2,109 cells). e, Associations between inferred APOE and CLU signalling to the TREM2-TYROBP receptor complex and predicted DAM acquisition probabilities, assessed using a spatial permutation test. f, Spatial distributions of homeostatic microglia, DAM and amyloid plaques in TauPS2APP and control mouse brain sections profiled using STARmap PLUS at 8 and 13 months. g, Inferred transitions between homeostatic microglia and DAM in TauPS2APP and control mice. h, Predicted probabilities of DAM acquisition (left, n = 637 cells) and transitions towards homeostatic-like states (right, n = 319 cells) as a function of distance to the nearest amyloid plaque, expressed in units of typical cell spacing. Statistical associations were assessed after accounting for cellular expression state. i, Volcano plot showing differential gene expression between DAM predicted to transition towards a homeostatic-like state and those predicted to retain DAM identity in TauPS2APP mice (P < 0.05 and |log₂ fold change| > 0.25; two-sided Mann-Whitney U test). In d,h, lines indicate binned means and shaded regions indicate bootstrap 95% confidence intervals.

Niche-FATE predicted transitions from homeostatic microglia to DAM predominantly in TgCRND8 mice (Fig. 5b). The estimated transition probability from homeostatic microglia to DAM increased from 5% between 2.5 and 5.7 months to 20% between 5.7 and 17.9 months, compared with 1-2% in wild-type mice sampled at 2.5, 5.7 and 13.4 months. These transitions are consistent with the progressive accumulation of DAM during amyloid pathology^44^.

At 5.7 months, homeostatic microglia predicted to acquire DAM identity expressed lower levels of homeostatic markers, including *P2ry12* and *Tmem119*, and higher levels of DAM markers, including *Trem2*, *Apoe* and *Cd68*, than those predicted to remain homeostatic (Fig. 5c). Predicted transition probabilities were higher near existing DAM and decreased with distance over approximately 100 µm (P = 1 x 10^−30^; Fig. 5d).

We next examined the niche signals associated with DAM acquisition (Fig. 5e; Supplementary Table 3). These included predicted interactions of APOE and clusterin with the TREM2-TYROBP receptor complex. The inferred strength of these interactions correlated positively with the probability of DAM acquisition between 2.5 and 5.7 months (r = 0.37 and 0.38, respectively) and between 5.7 and 17.9 months (r = 0.17 and 0.18, respectively; all P = 0.001). Both apolipoproteins accumulate around amyloid plaques and are known ligands of TREM2^47^, which promotes DAM acquisition^44,45^.

To examine whether these findings extended to a second disease model and spatial transcriptomics platform, we analysed STARmap PLUS data from TauPS2APP mice and controls at 8 and 13 months^13^. TauPS2APP mice develop both amyloid and tau pathology, which were mapped in the same tissue sections (Fig. 5f; Extended Data Fig. 7c-f). Niche-FATE again predicted a higher transition probability from homeostatic microglia to DAM in TauPS2APP mice (25%) than in controls (2%; Fig. 5g). Although plaque positions were not provided to Niche-FATE, homeostatic microglia closer to amyloid plaques had higher predicted probabilities of acquiring DAM identity (P = 4 x 10^−5^; Fig. 5h). This association was not significant when niche information was excluded (P = 0.55; Extended Data Fig. 7g).

Niche-FATE also assigned 32% of the transition probability from DAM at 8 months to homeostatic microglia at 13 months (Fig. 5g), suggesting that disease-associated identity may not be uniformly maintained during neurodegeneration. Unlike DAM acquisition, predicted transitions towards homeostasis were not significantly associated with plaque distance (P = 0.72; Fig. 5h). DAM predicted to transition towards homeostatic-like states expressed lower levels of DAM-associated and lysosomal genes, including *Trem2*, *Cd9*, *Apoe*, *Lyz2* and *Ctsb*, than those predicted to retain DAM identity (Fig. 5i; Supplementary Table 4). These findings suggest heterogeneity within the DAM population in its potential to retain disease-associated identity.

By linking predicted cellular transitions to local pathology and signalling environments, Niche-FATE generates hypotheses about microglial state plasticity that cannot be derived from changes in cellular abundance alone.

## Discussion

Niche-FATE integrates local signalling environments into temporal cell-state inference and prioritises candidate pathways associated with alternative predicted outcomes. Together, these capabilities extend spatial transcriptomic time-course analysis from reconstructing cellular trajectories to investigating their potential regulation by the tissue environment.

The value of niche information depends on what it adds beyond a cell’s intrinsic transcriptional state. In simulations, inference performance increased with the contribution of extrinsic signals to lineage choice. During organogenesis and digit tip regeneration, niche information helped distinguish transcriptionally related populations occupying distinct tissue environments. In AD, incorporating niche information had relatively little effect on aggregate transition estimates but helped resolve spatial variation in predicted DAM acquisition relative to amyloid plaques, whose positions were not supplied to the model. These findings highlight the complementary contributions of intrinsic transcriptional programmes and local environments to the inference of cellular dynamics.

Beyond improving temporal inference, Niche-FATE generates hypotheses about cellular plasticity that are difficult to resolve from state abundance alone. In AD, Niche-FATE predicted transitions from DAM towards homeostatic-like states despite ongoing pathology. Previous studies have demonstrated microglial state plasticity during development and remyelination^48,49^ and identified mechanisms regulating microglial states^49,50^. Whether the predicted DAM-to-homeostatic transitions occur during ongoing neurodegeneration could be investigated using state-specific genetic fate mapping.

Several limitations should be considered. First, as with other optimal transport-based approaches, Niche-FATE infers transitions between independently sampled cell populations rather than directly tracking individual cells. Predicted transitions would require validation, especially when distinguishing state conversion from proliferation or survival. Niche-based matching is particularly informative when local environmental characteristics are retained across time points, even when global tissue architecture changes. Second, Niche-FATE estimates intercellular signalling from ligand and receptor gene expression and spatial proximity, which does not directly measure ligand availability, receptor activation, or downstream pathway activity. Because these signalling scores also contribute to transport, their associations with inferred fates prioritise candidate regulators rather than independently establish regulatory relationships. Perturbation of candidate signalling pathways, combined with lineage tracing, will be necessary to establish functional contributions. Third, the representation of cellular niches depends on transcript detection and gene coverage. For example, the limited gene coverage constrained the identification of candidate signalling pathways associated with microglial state transitions. Higher-coverage spatial transcriptomics and the integration of spatial proteomic measurements could improve the representation of cellular signalling environments.

Niche-FATE provides a framework for connecting cellular dynamics to their tissue environments and translating spatial transcriptomic time courses into testable hypotheses about cell-state regulation.

## Methods

### Niche-FATE framework

#### Overview

Niche-FATE infers cell-state transitions and cell fates from spatiotemporal transcriptomics data. It matches cells of an earlier section to cells of a later section by optimal transport (OT). The matching cost combines the expression state of each cell with two descriptions of its niche: the ligand–receptor signals it receives and an embedding of its spatial neighbourhood. From the matching, Niche-FATE estimates state transitions and a fate probability for every cell. It then identifies the received signals associated with each fate.

#### Expression state

Counts were normalized to the total counts per cell and log-transformed. The expression state of each cell was represented by principal components computed jointly across sections.

#### Received signals

For each ligand-receptor pair from CellPhoneDB^23^, the signal received by cell *i* is computed as in CytoSignal^24^:

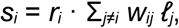

where *r_i_* is the receptor expression of cell *i*, *ℓ_j_* is the ligand expression of cell *j* and *w_ij_* weights cell *j* by its distance to cell *i*. For secreted ligands, *w_ij_* is a Gaussian function of distance; for contact-dependent ligands, it includes only direct neighbours. Scores are averaged within each signalling pathway. Distances are measured in units of the typical spacing between neighbouring cells.

#### Niche embedding

The expression of each cell’s neighbourhood is summarized by a graph autoencoder^51^. Each section is represented as a spatial nearest-neighbour graph, with the expression principal components of each cell as input features *H*^(0)^. Graph convolution layers encode each neighbourhood into a low-dimensional embedding:

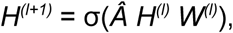

where *Â* is the row-normalized adjacency matrix, so that each layer averages features over neighbouring cells, *W*^(l)^ are learned weights and *σ* is a nonlinear activation. The input is reconstructed from the final embedding, and the model is trained to minimize the reconstruction error.

#### Matching

For a source cell *i* and a target cell *j*, the cost is

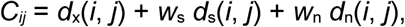

where *d*_x_, *d*_s_ and *d*_n_ are squared Euclidean distances in the expression state, received signals and niche embedding. Each term is scaled to a comparable range, and *w*_s_ and *w*_n_ set the weight of the niche module, which consists of the received signals and the niche embedding. The transport plan *P*\*, in which *P*\* is the mass moved from cell *i* to cell *j*, is

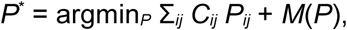

where *M*(*P*) is a Kullback–Leibler penalty on the difference between the mass that leaves or reaches each cell and its expected mass. Because this is a penalty rather than a hard constraint, cells can proliferate or die between time points. We solve this problem by entropic unbalanced OT^52^. When the tissue layout is largely conserved between sections, a fused Gromov-Wasserstein term on spatial distances within each section can be added, so that matched cells also keep their spatial relations^53^. We used this term for the fixed-layout simulations and for mouse organogenesis, where consecutive embryos share the same body plan. Matching can be restricted to selected cell groups, such as one lineage.

#### Fate probabilities

The fate probability of source cell *i* for target state *k* is the share of its transported mass that reaches cells of state *k*:

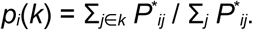

State transitions, shown as Sankey diagrams, are the mass flows between source and target states.

#### Signals associated with fate

Each received signal is tested for association with the fate probability among the source cells of interest. We compute the partial rank correlation between signal and fate probability, adjusting for the cell’s own expression state, local cell density and, optionally, the cell-type composition of the neighbourhood. P-values are obtained from 1,000 permutations of fate probabilities between spatial blocks, which keep the spatial structure of the data. The strength of the association is reported as a z-score relative to these permutations. When several sections are analysed, z-scores are combined across sections by Stouffer’s method.

### Simulated data

We simulated spatial time courses with two time points based on scMultiSim^25^. Each simulation contains a progenitor lineage that splits into two progeny states, A and B, and two sender lineages. It has 2,000 cells per time point, 1,000 genes and 20 ligand-receptor pairs. Each ligand is expressed mainly by one sender lineage or by progeny cells, so the signal a cell receives depends on its neighbours. The fate of each progenitor depends on the signals it receives through three driver pairs, and on its own expression. The other 17 pairs are decoys. At the second time point, every cell is replaced by a descendant: progenitors that have decided become progeny A or B, and other cells advance along their own lineage.

We simulated two tissue scenarios. In shifting niches, the second section has a new layout, and each cell is placed in a niche similar to that of its parent. In the fixed layout, the second section keeps the layout of the first, with small displacements. Each scenario has four signal strengths (none, weak, medium and strong), with five replicates each. Sequencing depth ranges from about 1,100 to 3,000 transcripts per cell.

### Evaluation

#### Simulations

Fate prediction was scored on progenitors that had decided by the second time point. For each method, the predicted probability of state B was compared with the true fate by the area under the receiver operating characteristic curve (AUROC). The best achievable AUROC was computed from the true fate probability. Signal recovery was scored by the area under the precision-recall curve (AUPRC) over the 20 ligand-receptor pairs, with the three driver pairs as positives. For every method, signals were ranked by the same association test applied to their fate probabilities. In the control without the niche module, cells were matched on expression alone. In the niche-shuffled control, cell positions were permuted within each section before the niche module was computed.

#### Mouse organogenesis

We used the curated list of expected cell-type transitions from the moscot study^17^. For each source cell type, accuracy is the share of its transported mass that reaches an expected descendant. We report the mean over cell types, together with the accuracy of a random matching.

### Compared methods

We compared Niche-FATE with the following methods. Fate probabilities were computed from each method’s cell matching in the same way as for Niche-FATE.

**WOT^14^** matches cells between time points by OT on expression, allowing cells to proliferate or die between time points through unbalanced OT. It does not use spatial information.

**moscot17** extends OT to spatial data by adding a fused Gromov-Wasserstein term on spatial coordinates. We used this spatiotemporal mode when each time point had one section, and its expression-only temporal mode for the digit data, where each time point pooled many sections.

**STORIES^18^** learns a potential over expression space from spatial time courses and predicts the future state of each cell. To obtain a matching, we transported the predicted states onto the observed cells of the next time point by OT on expression. Because STORIES uses spatial position within a section, we did not apply it to the digit data, where each time point pooled many sections.

**spaTrack^16^** infers transitions between time points by OT that combines expression and spatial position. Like moscot, it uses spatial position directly in the matching. We therefore did not apply spaTrack to the digit data.

### Datasets and analyses

#### Mouse organogenesis

We used Stereo-seq sections of mouse embryos from E9.5 to E16.5, with one sagittal section per day^11^. The data contain 518,165 spots of about 25 µm, each of which can contain several cells, and seven pairs of consecutive time points.

#### Mouse digit tip regeneration

We used a Xenium dataset of 179 mouse digits (about 510,000 cells) after distal amputation, which regenerates, or proximal amputation, which heals with a scar^36^. Regenerative digits were sampled at 5, 7, 10 and 14 days post amputation (dpa), and non-regenerative digits at 5, 7 and 14 dpa. At 5 dpa, signals were averaged per digit and compared between the two types of digits by Mann–Whitney U test.

#### Amyloid model TgCRND8

We used public Xenium data from 10x Genomics, with one coronal brain section each from TgCRND8 mice at 2.5, 5.7 and 17.9 months and wild-type mice at 2.5, 5.7 and 13.4 months^46^. The six sections contain 346,093 cells measured with a 347-gene panel. Amyloid plaques were imaged on the 17.9-month TgCRND8 section.

#### Amyloid and tau model TauPS2APP

We used STARmap PLUS data from TauPS2APP and control mice at 8 and 13 months, with two mice per group and one section per mouse^13^. The eight sections contain 72,165 cells measured for 2,766 genes. Amyloid plaques and phosphorylated tau were imaged in the same sections. The association between fate probability and distance to the nearest plaque was tested as for signals, in each mouse, and combined across mice. DAM predicted to revert were compared with DAM predicted to remain by Mann-Whitney U test for each gene.

### Statistics

All tests were two-sided. Unless stated otherwise, n is the number of cells; in digit-level comparisons, n is the number of digits. Binned plots show means with bootstrap 95% confidence intervals.

## Data availability

The mouse organogenesis data are available from the MOSTA database^11^. The digit tip data are available from GEO under accession GSE326776. The TgCRND8 Xenium data are available from 10x Genomics. The STARmap PLUS data are available at https://singlecell.broadinstitute.org/single_cell/study/SCP1375. Simulation datasets are available at https://doi.org/10.5281/zenodo.24346489.

## Code availability

The Niche-FATE package and tutorials are available at https://github.com/jiang-junyao/Niche-FATE.

## Author contributions

J.J. and D.W.K. conceived the study. J.J. developed Niche-FATE, implemented the software and performed the analyses. D.W.K. supervised the study and acquired funding. J.J. and D.W.K. wrote the manuscript.

---

**Extended Data Fig. 1.**
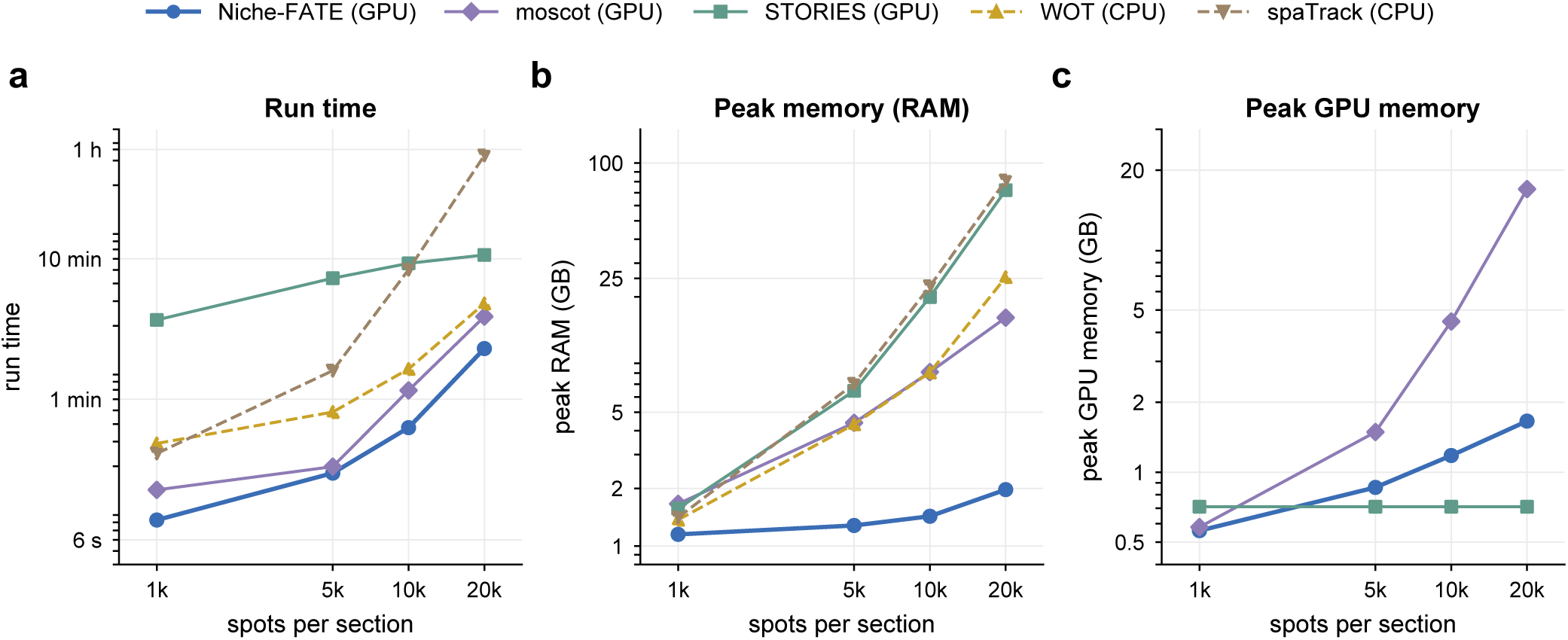
Computational runtime and memory benchmarks. Methods were evaluated using datasets containing 1,000, 5,000, 10,000 and 20,000 spatial spots per section. **a,** Runtime. **b,** Peak RAM usage. **c,** Peak GPU memory usage. GPU-based methods were run on a single NVIDIA L40S GPU, and all methods used 16 CPU cores.

**Extended Data Fig. 2.**
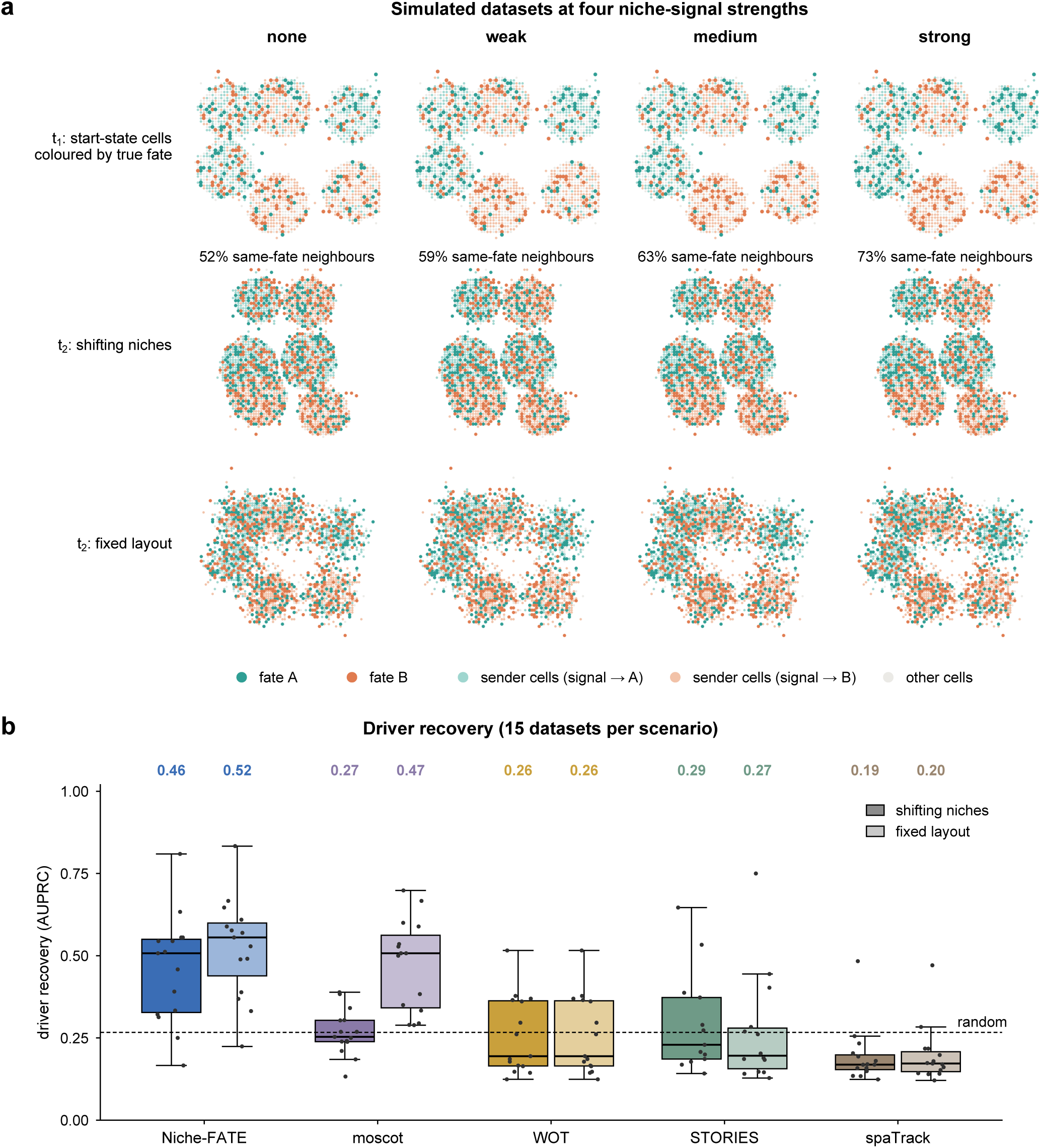
Simulated datasets and recovery of fate-determining signals. **a,** Representative simulated datasets at four niche signal strengths. The top row shows progenitors at the initial time point coloured by their true subsequent fates. Percentages indicate the proportion of neighbouring progenitors sharing the same fate. The middle and bottom rows show the subsequent time point in tissues with shifting niches and a fixed layout, respectively. **b,** Recovery of known fate-determining signals, measured by the area under the precision–recall curve (AUPRC), in tissues with shifting niches (dark) and a fixed layout (light). Each simulated dataset contains three fate-determining signals among 20 candidate signals. Signals were ranked using the same association test for all methods. n = 15 simulated datasets per configuration. Numbers above the box plots indicate mean AUPRC values. The dashed line indicates the random ranking baseline.

**Extended Data Fig. 3.**
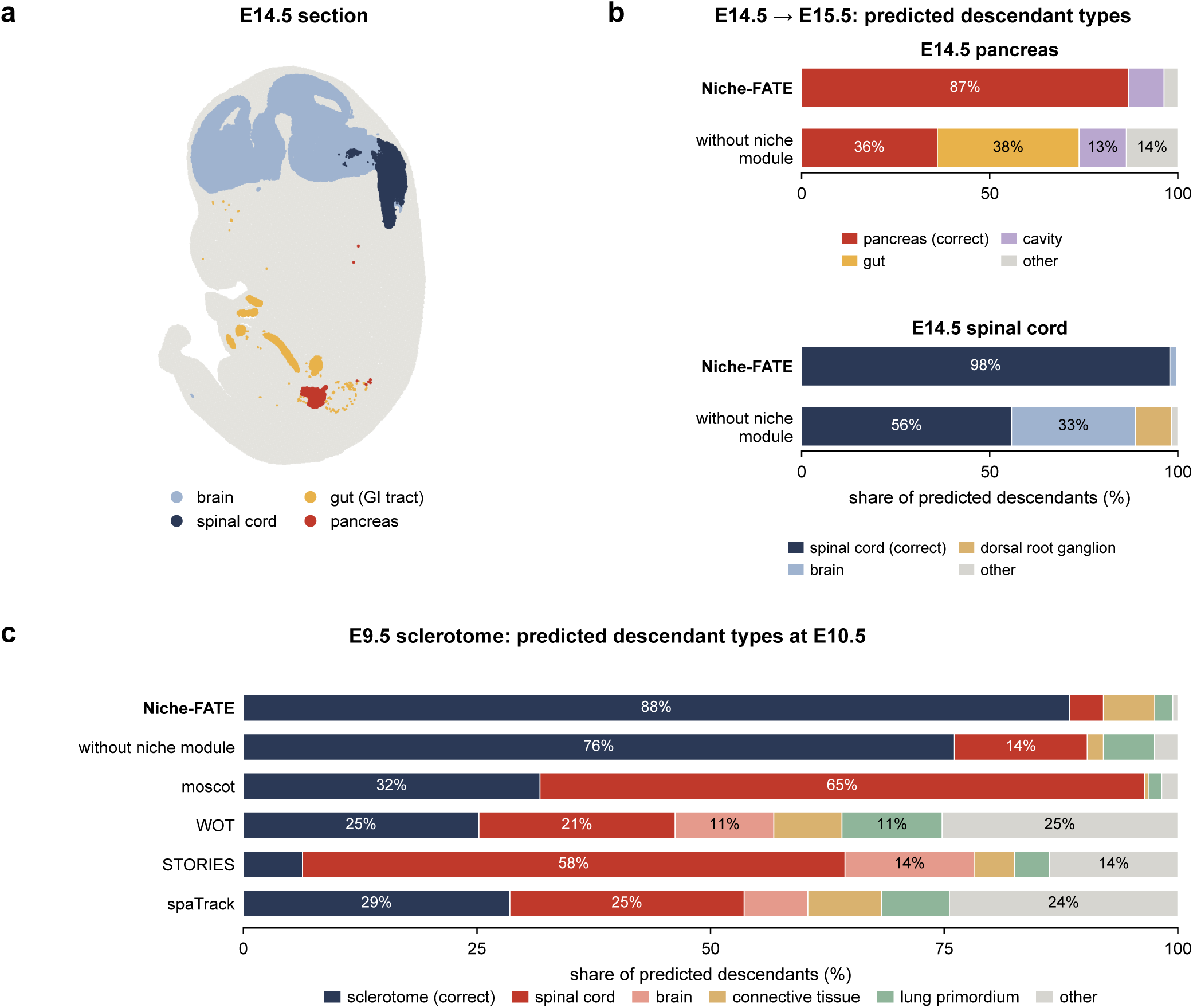
The niche module reduces developmentally inconsistent transitions during mouse organogenesis. **a,** E14.5 embryonic section showing the anatomical locations of the pancreas, gut, spinal cord and brain. **b,** Predicted E15.5 descendant populations of E14.5 pancreas (top) and spinal cord (bottom), inferred with and without the niche module. **c,** Comparison of predicted E10.5 descendant populations of E9.5 sclerotome across different methods.

**Extended Data Fig. 4.**
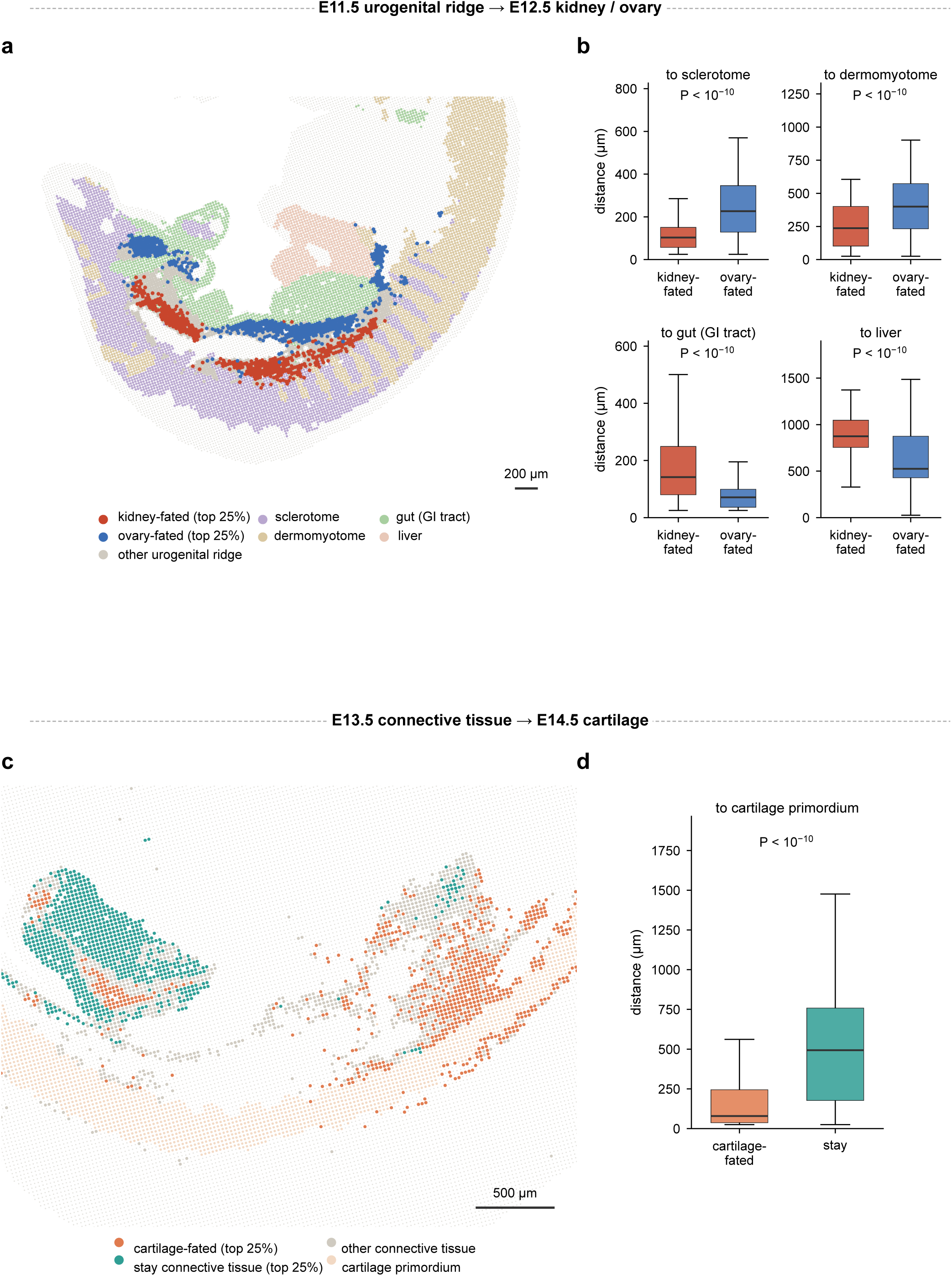
Predicted developmental outcomes are associated with distinct anatomical environments. **a,** Spatial distribution of E11.5 urogenital ridge spots with the highest predicted probabilities of transitioning towards kidney (red, top 25%) or ovarian (blue, top 25%) identity. **b,** Distances from these populations to the sclerotome, dermomyotome, gut and liver. **c,** Spatial distribution of E13.5 connective tissue spots with the highest predicted probabilities of acquiring cartilage identity (orange, top 25%) or retaining connective tissue identity (green, top 25%). **d,** Distances from these populations to the cartilage primordium. P values in **b,d** were calculated using two-sided Mann–Whitney U tests. Scale bars, 200 µm (**a**) and 500 µm (**c**).

**Extended Data Fig. 5.**
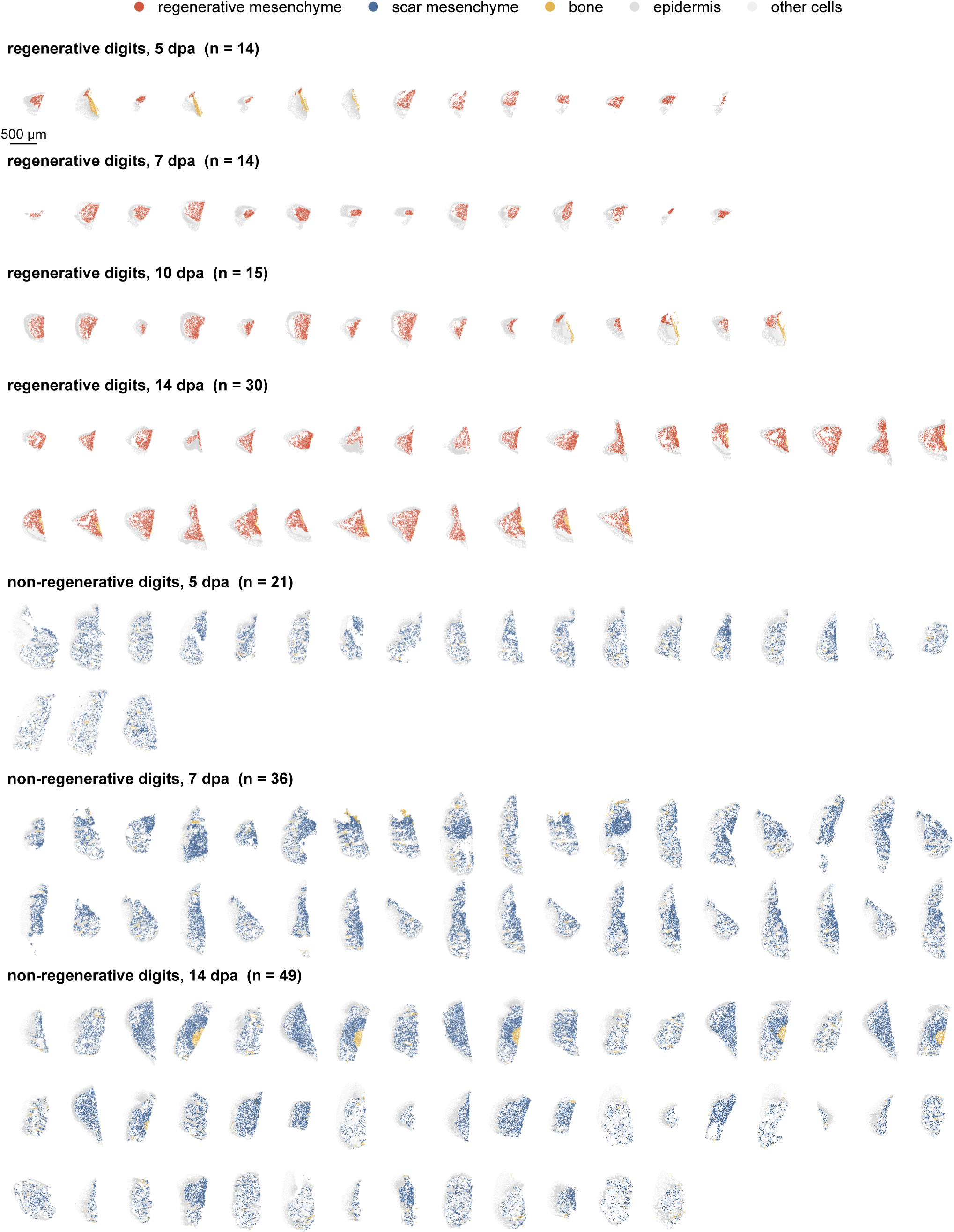
Overview of the mouse digit tip regeneration dataset. Spatial transcriptomic sections grouped by amputation outcome and time point, displayed at the same spatial scale. Colours indicate annotated cell populations, as in Fig. 4b. n indicates the number of digits. Scale bar, 500 µm.

**Extended Data Fig. 6.**
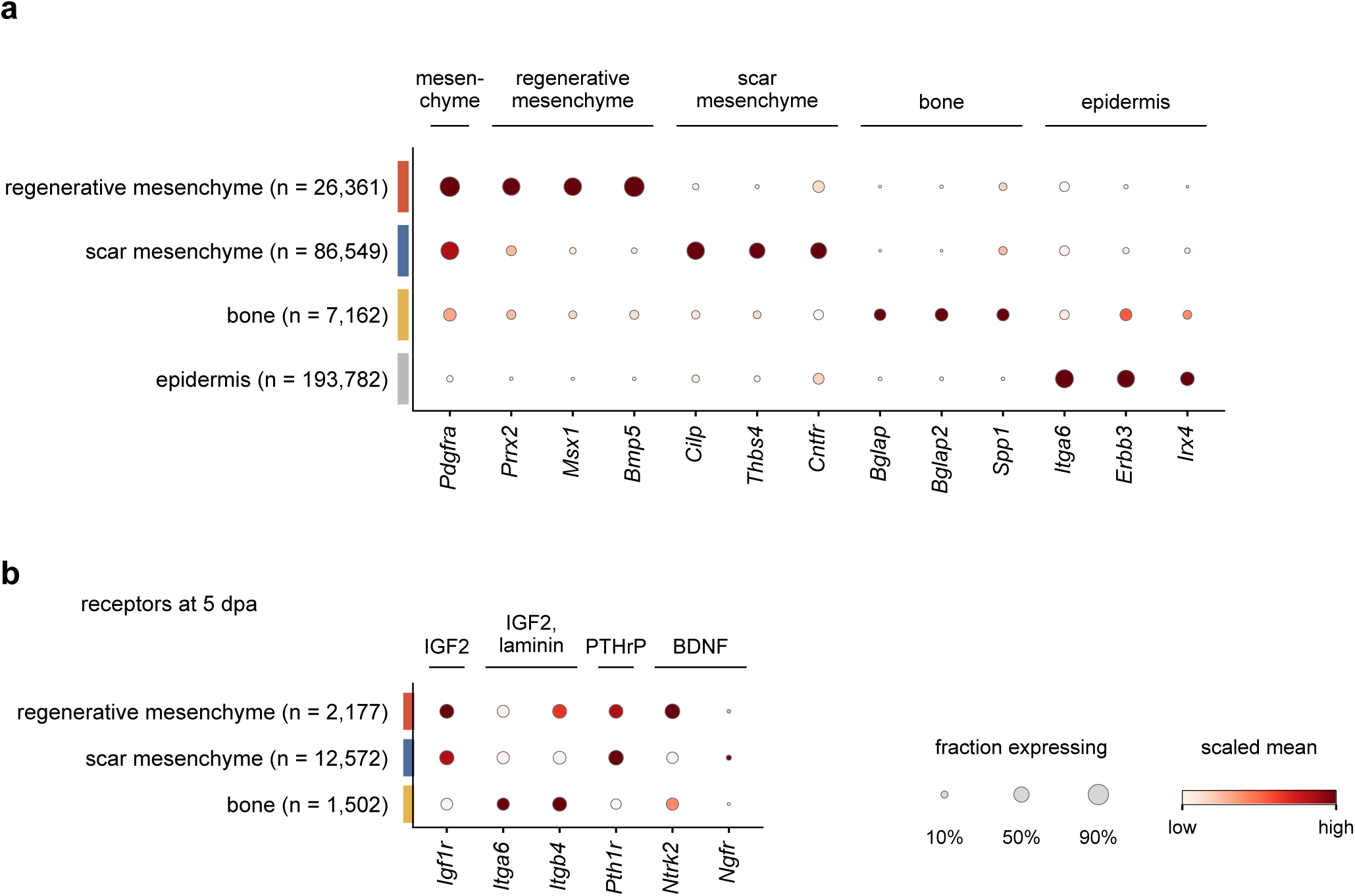
Expression of cell-type markers and signalling receptors during digit tip regeneration. **a,** Expression of representative marker genes across annotated cell populations. **b,** Expression of receptors corresponding to the niche signals identified in Fig. 4d at 5 dpa. Dot size indicates the proportion of cells expressing each gene, and colour indicates scaled mean expression. n indicates the number of cells.

**Extended Data Fig. 7.**
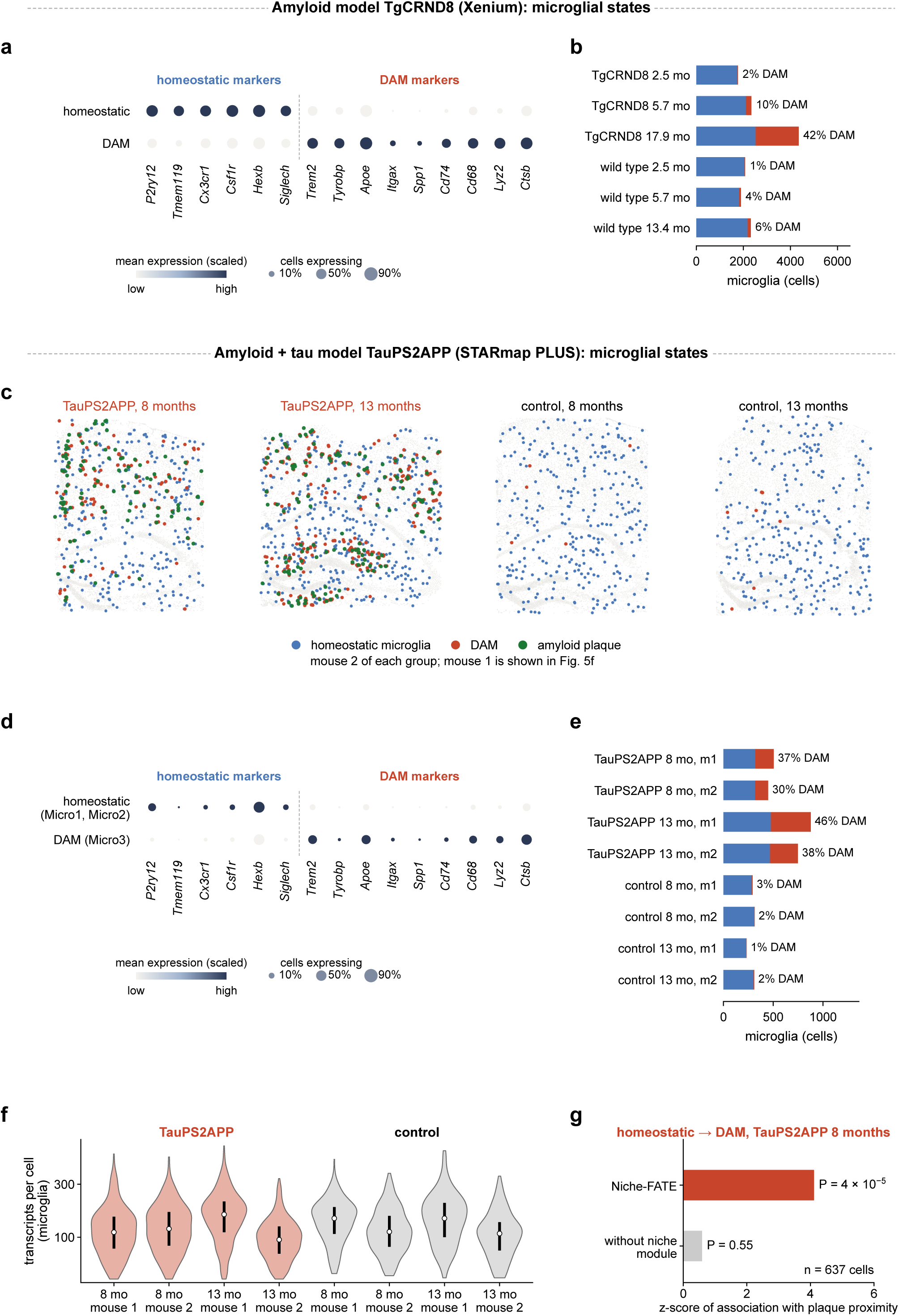
Overview of Alzheimer’s disease datasets and the contribution of niche information to predicted DAM acquisition. **a,** Expression of homeostatic and DAM marker genes in the TgCRND8 Xenium dataset. **b,** Numbers of microglia in TgCRND8 and wild-type mouse brain sections. Percentages indicate the proportion of DAM. **c,** Spatial distributions of microglial populations and amyloid plaques in sections from the second mouse in each TauPS2APP and control group, corresponding to Fig. 5f. **d,** Expression of homeostatic and DAM marker genes in the TauPS2APP STARmap PLUS dataset. **e,** Numbers of microglia in TauPS2APP and control mouse brain sections. Percentages indicate the proportion of DAM. **f,** Distribution of detected transcripts per microglial cell in individual TauPS2APP and control mouse brain sections. **g,** Association between predicted DAM acquisition probability and amyloid plaque proximity in homeostatic microglia from 8-month-old TauPS2APP mice, with and without the niche module (n = 637 cells). Bars indicate association z-scores calculated after accounting for cellular expression state. In **a,d**, dot size indicates the proportion of cells expressing each gene, and colour indicates scaled mean expression.

## References

1. Martin, B. L. & Kimelman, D. Canonical Wnt signaling dynamically controls multiple stem cell fate decisions during vertebrate body formation. Dev. Cell 22, 223–232 (2012).

2. Gouti, M. et al. In vitro generation of neuromesodermal progenitors reveals distinct roles for Wnt signalling in the specification of spinal cord and paraxial mesoderm identity. PLoS Biol. 12, e1001937 (2014).

3. Row, R. H., Tsotras, S. R., Goto, H. & Martin, B. L. The zebrafish tailbud contains two independent populations of midline progenitor cells that maintain long-term germ layer plasticity and differentiate in response to local signaling cues. Development 143, 244–254 (2016).

4. Yuan, Y. et al. 3D reconstruction of a human Carnegie stage 9 embryo provides a snapshot of early body plan formation. Cell Stem Cell 32, 1006–1024 (2025).

5. Öhlund, D. et al. Distinct populations of inflammatory fibroblasts and myofibroblasts in pancreatic cancer. J. Exp. Med. 214, 579–596 (2017).

6. Biffi, G. et al. IL1-induced JAK/STAT signaling is antagonized by TGFβ to shape CAF heterogeneity in pancreatic ductal adenocarcinoma. Cancer Discov. 9, 282–301 (2019).

7. Jones, M. G. et al. Spatiotemporal lineage tracing reveals the dynamic spatial architecture of tumor growth and metastasis. Nat. Genet. 58, 2398–2410 (2026).

8. Weinreb, C., Rodriguez-Fraticelli, A., Camargo, F. D. & Klein, A. M. Lineage tracing on transcriptional landscapes links state to fate during differentiation. Science 367, eaaw3381 (2020).

9. Jain, S. & Eadon, M. T. Spatial transcriptomics in health and disease. Nat. Rev. Nephrol. 20, 659–671 (2024).

10. Ståhl, P. L. et al. Visualization and analysis of gene expression in tissue sections by spatial transcriptomics. Science 353, 78–82 (2016).

11. Chen, A. et al. Spatiotemporal transcriptomic atlas of mouse organogenesis using DNA nanoball-patterned arrays. Cell 185, 1777–1792 (2022).

12. Wei, X. et al. Single-cell Stereo-seq reveals induced progenitor cells involved in axolotl brain regeneration. Science 377, eabp9444 (2022).

13. Zeng, H. et al. Integrative in situ mapping of single-cell transcriptional states and tissue histopathology in a mouse model of Alzheimer’s disease. Nat. Neurosci. 26, 430–446 (2023).

14. Schiebinger, G. et al. Optimal-transport analysis of single-cell gene expression identifies developmental trajectories in reprogramming. Cell 176, 928–943 (2019).

15. Sha, Y., Qiu, Y., Zhou, P. & Nie, Q. Reconstructing growth and dynamic trajectories from single-cell transcriptomics data. Nat. Mach. Intell. 6, 25–39 (2024).

16. Shen, X. et al. Inferring cell trajectories of spatial transcriptomics via optimal transport analysis. Cell Syst. 16, 101194 (2025).

17. Klein, D. et al. Mapping cells through time and space with moscot. Nature 638, 1065–1075 (2025).

18. Huizing, G. J. et al. STORIES: learning cell fate landscapes from spatial transcriptomics using optimal transport. Nat. Methods 23, 522–531 (2026).

19. Peng, Q., Zhou, P. & Li, T. stVCR: spatiotemporal dynamics of single cells. Nat. Methods 23, 542–553 (2026).

20. Bryan, J. P., Farhi, S. L. & Cleary, B. Accurate trajectory inference in time-series spatial transcriptomics with structurally-constrained optimal transport. Nat. Commun. 17, 8075 (2026).

21. Gu, Y. et al. Topological velocity inference from spatial transcriptomic data. Nat. Biotechnol. 44, 771–782 (2026).

22. Yan, L., Zhang, D. & Sun, X. Decoding cell state transitions driven by dynamic cell–cell communication in spatial transcriptomics. Nat. Comput. Sci. 6, 193–207 (2026).

23. Troulé, K. et al. CellPhoneDB v5: inferring cell–cell communication from single-cell multiomics data. Nat. Protoc. 20, 3412–3440 (2025).

24. Liu, J. et al. CytoSignal detects locations and dynamics of ligand–receptor signaling at cellular resolution from spatial transcriptomic data. Nat. Genet. 58, 1396–1408 (2026).

25. Li, H., Zhang, Z., Squires, M., Chen, X. & Zhang, X. scMultiSim: simulation of single-cell multi-omics and spatial data guided by gene regulatory networks and cell–cell interactions. Nat. Methods 22, 982–993 (2025).

26. Kicheva, A. & Briscoe, J. Control of tissue development by morphogens. Annu. Rev. Cell Dev. Biol. 39, 91–121 (2023).

27. Bunce, C., McKey, J. & Capel, B. Concerted morphogenesis of genital ridges and nephric ducts in the mouse captured through whole-embryo imaging. Development 148, dev199208 (2021).

28. Chassot, A. A. et al. Activation of beta-catenin signaling by Rspo1 controls differentiation of the mammalian ovary. Hum. Mol. Genet. 17, 1264–1277 (2008).

29. Tomizuka, K. et al. R-spondin1 plays an essential role in ovarian development through positively regulating Wnt-4 signaling. Hum. Mol. Genet. 17, 1278–1291 (2008).

30. Matsubara, N., et al. A receptor tyrosine kinase, Sky, and its ligand Gas 6 are expressed in gonads and support primordial germ cell growth or survival in culture. Dev. Biol. 180, 499–510 (1996).

31. Matsushita, Y. et al. Hes1 marks peri-condensation mesenchymal cells that generate both chondrocytes and perichondrial cells in early bone development. J. Biol. Chem. 299, 104805 (2023).

32. Day, T. F., Guo, X., Garrett-Beal, L. & Yang, Y. Wnt/beta-catenin signaling in mesenchymal progenitors controls osteoblast and chondrocyte differentiation during vertebrate skeletogenesis. Dev. Cell 8, 739–750 (2005).

33. Hill, T. P., Später, D., Taketo, M. M., Birchmeier, W. & Hartmann, C. Canonical Wnt/beta-catenin signaling prevents osteoblasts from differentiating into chondrocytes. Dev. Cell 8, 727–738 (2005).

34. Lebensohn, A. M. & Rohatgi, R. R-spondins can potentiate WNT signaling without LGRs. eLife 7, e33126 (2018).

35. Takeo, M. et al. Wnt activation in nail epithelium couples nail growth to digit regeneration. Nature 499, 228–232 (2013).

36. Purushothaman, S. et al. The nail mesenchyme creates a regeneration-specific ligand environment that orchestrates mammalian regeneration versus fibrotic wound healing. Preprint at bioRxiv 10.64898/2026.04.18.719303 (2026).

37. Huang, Y. et al. Igf signaling is required for cardiomyocyte proliferation during zebrafish heart development and regeneration. PLoS ONE 8, e67266 (2013).

38. Shen, H. et al. Mononuclear diploid cardiomyocytes support neonatal mouse heart regeneration in response to paracrine IGF2 signaling. eLife 9, e53071 (2020).

39. Nornberg, B. F., Almeida, D. V., Figueiredo, M. A. & Marins, L. F. GH indirectly enhances the regeneration of transgenic zebrafish fins through IGF2a and IGF2b. Transgenic Res. 25, 743–749 (2016).

40. Chen, C.-H. et al. Transient laminin beta 1a induction defines the wound epidermis during zebrafish fin regeneration. PLoS Genet. 11, e1005437 (2015).

41. Chen, Z.-L. & Strickland, S. Laminin γ1 is critical for Schwann cell differentiation, axon myelination, and regeneration in the peripheral nerve. J. Cell Biol. 163, 889–899 (2003).

42. Wang, M., Nasiri, A. R., Broadus, A. E. & Tommasini, S. M. Periosteal PTHrP regulates cortical bone remodeling during fracture healing. Bone 81, 104–111 (2015).

43. Clow, C. & Jasmin, B. J. Brain-derived neurotrophic factor regulates satellite cell differentiation and skeletal muscle regeneration. Mol. Biol. Cell 21, 2182–2190 (2010).

44. Keren-Shaul, H. et al. A unique microglia type associated with restricting development of Alzheimer’s disease. Cell 169, 1276–1290 (2017).

45. Krasemann, S. et al. The TREM2-APOE pathway drives the transcriptional phenotype of dysfunctional microglia in neurodegenerative diseases. Immunity 47, 566–581 (2017).

46. 10x Genomics. Xenium In Situ analysis of Alzheimer’s disease mouse model brain coronal sections from one hemisphere over a time course. https://www.10xgenomics.com/datasets/xenium-in-situ-analysis-of-alzheimers-disease-mouse-model-brain-coronal-sections-from-one-hemisphere-over-a-time-course-1-standard (2023).

47. Yeh, F. L., Wang, Y., Tom, I., Gonzalez, L. C. & Sheng, M. TREM2 binds to apolipoproteins, including APOE and CLU/APOJ, and thereby facilitates uptake of amyloid-beta by microglia. Neuron 91, 328–340 (2016).

48. Barclay, K. M. et al. An inducible genetic tool to track and manipulate specific microglial states reveals their plasticity and roles in remyelination. Immunity 57, 1394–1412 (2024).

49. Hamagami, N. et al. State-specific enhancer landscapes govern microglial plasticity. Immunity 59, 288–305 (2026).

50. Ayata, P. et al. Lymphoid gene expression supports neuroprotective microglia function. Nature 648, 157–165 (2025).

51. Kipf, T. N. & Welling, M. Semi-supervised classification with graph convolutional networks. In Proc. 5th International Conference on Learning Representations (ICLR, 2017).

52. Cuturi, M. Sinkhorn distances: lightspeed computation of optimal transport. Adv. Neural Inf. Process. Syst. 26, 2292–2300 (2013).

53. Vayer, T., Courty, N., Tavenard, R., Chapel, L. & Flamary, R. Optimal transport for structured data with application on graphs. In Proc. 36th International Conference on Machine Learning Vol. 97, 6275–6284 (PMLR, 2019).

